# *Butyrivibrio azoria* sp. nov., a novel cellulolytic microorganism isolated from the rumen of a Holstein dairy cow

**DOI:** 10.64898/2026.07.16.739024

**Authors:** Kayla Calapa, Jordan Embree, Rachel Bock, Jolie LoBrutto, Mallory Embree

## Abstract

This study describes the characteristics of NATIVEDY162^T^ (= JL13D10^T^ = NRRL B-68525^T^) a novel bacterium isolated from the rumen of a healthy Holstein dairy cow. NATIVEDY162^T^ was discovered to be an obligately anaerobic, slightly curved, rod that stains Gram-negative, and presents as singlets or short chains. Tests confirmed it is indole-, catalase-, oxidase-negative, and it is not motile. NATIVEDY162^T^ indicated a growth preference within the pH range of 6.5-7.5 with optimal growth at pH 7.0. Carbon panel assays found NATIVEDY162^T^ can utilize D-glucose, L-rhamnose, esculin/ferric citrate, D-lactose, and D-trehalose, whereas weak growth was observed on D-xylose and D-cellobiose. It was also capable of degrading starch and digesting both soluble and insoluble forms of cellulose, with genomic analysis providing further support by revealing a diverse array of carbohydrate-active enzymes (CAZymes) targeting starch and plant structural components like pectin and cellulose. Fermentation of glucose by NATIVEDY162^T^ resulted in the major fermentation products lactate and butyrate. Phylogenetic analysis of the 16S rRNA gene positioned NATIVEDY162^T^ in close relation to other *Butyrivibrio* species. Genome comparisons using BLAST ANI identified its closest relative as *Butyrivibrio proteoclasticus* (75.7% identity); however, the similarity did not meet the 95% threshold for species identification. Phylogenetic, genomic, and chemotaxonomic findings highlight differences between NATIVEDY162^T^ and other *Butyrivibrio* species, indicating it is a novel species. NATIVEDY162^T^ was isolated from a lactating Holstein dairy cow located in California’s San Joaquin Valley, a region rich in Portuguese influence from Azorean migrants who played a key role in the development of the California dairy industry, producing most of the state’s milk by the mid-20th century. Given this historical context, we propose the name *Butyrivibrio azoria* type strain NATIVEDY162^T^ in honor of the significant contributions of Azorean and Portuguese dairy farmers in the region.

---

The bovine ruminal microbiome is established at birth and undergoes dynamic transformations throughout growth from calf to adulthood, with shifting populations of various types of bacteria, fungi, and phage shaping its digestive system [1-3]. The microbiome is intertwined with the ruminant’s health, digestion, nutrient absorption potential, overall production efficiency, and milk quality, and is influenced by diet, health events, changes in weather, dysbiotic episodes, and feed additives throughout the ruminant’s life [1, 4-10]. Bovine feed is abundant in plant matter such as grain, straw, pulp, and silage structurally made up of cellulose, hemicellulose, pectin, and lignin [8, 11]. Since cattle cannot digest these components independently, the rumen microbiome plays an indispensable, synergistic role in breaking them down within the rumen [6, 8, 12-13].

Microorganisms employ diverse strategies to colonize and degrade plant matter, including bacterial quorum sensing, pili attachment, and production of extracellular polymeric substances (EPS) [14-15]. Once attached to plant substrates, initial microbial colonizers degrade surficial layers, which allows for subsequent microbes to further breakdown inner layers as they become exposed [1, 13]. Sugars released from these fibers can be fermented into volatile fatty acids that are absorbed through the gastrointestinal tract to provide energy to the animal [12, 16-17].

*Butyrivibrio* are ubiquitous in ruminants and rank among the most abundant genera in the rumen of dairy cattle, alongside *Prevotella, Pseudobutyrivibrio*, and *Ruminococcus* [6, 18-19]. Many *Butyrivibrio* produce enzymes that assist in the breakdown of hemicellulose and the structural matrix of plant matter in bovine feed [1, 7, 18, 20-22]. The type strain *Butryrivibrio fibrisolvens* D1^T^, for example, was first isolated in 1956 by Bryant and Small while culturing microbes from cattle rumen and was determined to have the ability to break down plant cell wall components pectin and xylan [23]. The genus *Butyrivibrio* currently has three identified species to date: *B. fibrisolvens, B. hungatei*, and *B. proteoclastic*us [23, 24]. *B. crossotus* was recently re-classified to *Eshraghiella crossota* [25-26].

*Butyrivibrio spp*. are described as obligately anaerobic short, curved, usually flagellated rods that can have variable motility. They Gram stain negative but have a membrane structure more similar to Gram-positive microorganisms [27, 28]. In addition to being isolated from rumen fluid, they have also been found in human, rabbit, and horse fecal samples [27]. The name *Butyrivibrio* can be broken into ‘butyricus’ and ‘vibrio,’ which is translated from Latin as ‘a butyric vibrio,’ a label indicative of their curved rod shape and characteristic butyric acid production [24].

*Butyrivibrio* are polyphyletic and tend to cluster with *Clostridium* subphylum XIVa [28, 29-30]. Palevich et al. 2019 determined there are phylogenetically three distinct clusters of *Butyrivibrio* species, which are somewhat predictive of their degradative capabilities: cluster one which includes *B. proteoclasticus* and *B. hungatei* had the most narrow substrate utilization profile, cluster two strains made up of unidentified *Butyrivibrio spp*. all exhibited consistent xylose utilization and an inability to grow on pectin, and a third *B. fibrisolvens* cluster which has similar capabilities of the first group and are capable of degrading pectin, xylan, and xylose [30-31]. Close neighbors include *Clostridium aminophilum, Agathobacter rectalis* (formerly *Eubacterium rectale*), and the genera *Pseudobutyrivibrio* and *Roseburia* [29, 31-32].

In this study, we investigate the phenotypic and genomic characteristics of NATIVEDY162^T^, a bacterium isolated from the rumen of a Holstein cow housed at a commercial dairy farm in Corcoran, CA, U.S.A.

## Isolation and Phenotypic Characterization

NATIVEDY162^T^ was sourced from the rumen of a Holstein dairy cow located at commercial dairy farm in Corcoran, CA. Initial growth and isolation was carried out using an unmodified anaerobic tryptic soy broth media (TSB; BD Biosciences, Franklin Lakes, NJ). Cells were cultured in this medium in an anaerobic environment (5% H_2_, 20% CO_2_, 75% N_2_) for a period of 72 hours at 37°C. When grown in a liquid medium, cells clump together to form aggregates. Cell morphological visualization was captured using an Olympus BX43 light microscope with an Excelis MPX-5C Pro camera attachment and CaptaVision v2.3.1.0 program. NATIVEDY162^T^ was viewed under 1000X magnification and presented as slightly curved rods (0.550-0.7 μm x 2.5-3.6 μm) found in singlets or short chains (Supplementary Figure 1). Gram stains were performed as outlined in the protocol by Beveridge, which confirmed NATIVEDY162^T^ stains Gram negative [33] (Supplementary Figure 2). When streaked for isolation on unmodified TSB agar solid media, colonies appear as opaque white, small, and circular with smooth margins.

### Genomic Characterization

NATIVEDY162^T^ total DNA was extracted from pure culture using a DNeasy PowerSoil Pro Kit (Qiagen, Germantown, MD). Short-read libraries were constructed with a KAPA HyperPlus kit (Roche, Indianapolis, IN) and steps carried out as instructed by the manufacturer’s protocol. Libraries were sequenced (1×300bp) on an Illumina MiSeq and sequencing adapters, indexes, and primers were trimmed using tool TrimGalore v0.6.4 [34].

In preparation for long read sequencing, high molecular weight DNA was extracted using a Monarch High Molecular Weight DNA Extraction Kit (New England BioLabs, Ipswich, MA). Long read libraries were processed with an Oxford Nanopore Technologies (ONT) Ligation v14 Sequencing Kit #SQK-LSK114, loaded onto a FLO-MIN114 flow cell, and sequenced with a MinION Mk1B device (ONT, Lexington, MA). Reads were basecalled with the ONT basecaller Dorado v0.4.0 and adapters subsequently removed with Porechop v0.5.1 [35-36]. Long reads smaller than 3500 bp and/or under a Q score threshold of 10 were quality filtered and checked with Filtlong v0.2.1 and Nanoplot v1.41.6, respectively [38-39]. A consensus assembly was created by combining quality filtered long and short reads using Unicycler v 0.5.0 and an assembly image was rendered with Bandage v0.8.1 [40-41].

Two complete, circular contigs were produced from the Unicycler hybrid assembly, both averaging greater than 1000X depth for long reads and greater than 38X for short reads as confirmed by Bedtools v2.26.0 [42]. Contig 1 was deposited to the National Center for Biotechnology Information (NCBI) GenBank under accession #CP172573 and contains 3,687,363 bp and 40.42 mol% G-C content. Contig 2 was deposited under #CP172574 and consists of 449,257 bp and 39.81 mol% G-C content. Table 1 outlines the genome size and G-C mol% differences between all known *Butyrivibrio, E. crossota* (formerly *B. crossotus*), and both identified *Pseudobutyrivibrio* species [5, 14, 20, 23-24, 26-27, 29, 31-32, 43].

**Table 1.** NATIVEDY162^T^ genomic, carbohydrate utilization, and fermentative characteristics compared to members of the genera *Butyrivibrio* and *Pseudobutyrivibrio*. Congo red agar and starch utilization assays were adapted from Sydlowski et al., 2015 and Lal, 2012 [64, 65], respectively, and as outlined in the Physiology and Chemotaxonomy section. All other strain’s chemotaxonomic and genomic traits were compiled from prior reports [5, 14, 20, 23-24, 26-27, 29, 31-32, 43].

| Characteristic | NATIVEDY162<sup>T</sup> | *E. crossota*<br>(*B. crossotus*)<br>ATCC 29175<sup>T</sup>/DSM 2876<sup>T</sup> | *B. fibrisolvens*<br>ATCC 19171<sup>T</sup>/D1<sup>T</sup> | *B. hungatei*<br>JK615<sup>T</sup> | *B. proteoclasticus*<br>B316<sup>T</sup> | *P. xylanivorans*<br>DSM 14809<sup>T</sup> | *P. ruminis*<br>DSM 9787<sup>T</sup> |
| --- | --- | --- | --- | --- | --- | --- | --- |
| Genome GC Content (%) | 40.4 | 37.0 | 39.5 | 44.8 | 40.0 | 42.1 | 40.5 |
| Genome Size (Mbp) | 4.14 | 2.52 | 4.93 | 3.39 | 4.40 | 3.42 | 3.02 |
| Major Fermentation Products (s) | A,B,L | A,B,F,L | B,F,L | A,B,F | B, F | B,F L,S | A,B,F,L |
| Able to Ferment: |  |  |  |  |  |  |  |
| Amygdalin | - | - | nd | - | + | nd | nd |
| Arabinose | w | - | nd | + | + | nd | + |
| Cellobiose | w | - | + | + | + | + | + |
| Cellulose | + | nd | nd | - | - | nd | nd |
| Esculin | + | - | + | + | + | + | nd |
| Fructose | - | w | + | - | + | nd | + |
| Glucose | + | w | + | + | + | + | + |
| Lactose | + | v | + | + | + | + | + |
| Mannitol | - | - | - | - | + | - | - |
| Starch | + | + | + | - | + | nd | - |
| Trehalose | + | - | nd | - | + | - | + |

Symbols: nd, no data; w, weak reaction; v, variable reaction; A, acetate; B, butyrate; F, formate; L, lactate; S, succinate.

Both contigs were further interrogated using the Bacterial and Viral Bioinformatics Resource Center (BV-BRC) v3.43.3 annotation tool which utilizes the Rapid Annotation using Subsystem Technology tool kit (RASTtk) pipeline [44-45]. Genome features show the second contig is characteristic of a secondary chromosome also known as a chromid. Chromids are estimated to be present in one in ten bacteria but tend to be more prevalent in specific genera (e.g. *Rhizobium, Afrobacterium*, and *Burkholderia*) and Gram-negative cells [20, 46-47]. Chromids are larger than a plasmid yet smaller than the primary chromosome in size, have a replication mechanism which is similar to plasmids (compared to the *oriC*/*dnaA* encoded replication instructions of the main chromosome), are within 1% G-C content of the main chromosome, and contain genes for cellular maintenance and metabolism [20, 46].

Contig two of NATIVEDY162^T^ has G-C content that is within 1 mol% of the main chromosome. It contains a ParA partitioning protein that matches closely with an unclassified *Butyrivbrio sp*. plasmid replication initiation protein when BLASTed in NCBI’s database (GenBank accession #WP_026511932). This second chromosome has genes which assist in metabolic functions of the cell involving starch, sucrose, and mannose utilization, and seems to have the ability to produce enzymes in which breakdown plant fibers (e.g. xylanases, endoglucanase, cellulase, glycosyl hydrolase). These functions resemble those identified on the chromid of *B. hungatei* MB2003 and align more closely with genes found on chromids rather than nonessential accessory genes located on plasmids and megaplasmids [20]. Consequently, the second complete contig of NATIVEDY162^T^ is presumed to be a chromid.

The main chromosome and chromid were further computationally analyzed to confirm the presence of carbohydrate active enzymes (CAZymes). Protein sequences were processed through the run_dbCAN v5.2.9 automatic CAZyme annotation pipeline using HMMER Hidden Markov Model for sequence alignment and homolog detection [48-50]. Cutoff thresholds were set to <1e-15 e-value and at least 70% coverage of the query protein sequence [48-50]. NATIVEDY162^T^ alignments resulted in a total of 219 hits, 81 of which included associated substrate specificity. Substrate mapping results indicated NATIVEDY162^T^ has a predicted substrate profile associated with the degradation of starch/glycogen (i.e. α-glucans) and plant matter (i.e. hits for xylan, arabinan, arabinogalactan, cellulose, β-glucans, galactans). A full list of CAZyme subfamily annotation and predicted substrate target results can be seen in Supplementary Table 2.

The 16S rRNA gene sequence was extracted from the genome assembly and first queried through NCBI’s Basic Local Alignment Search Tool (BLAST) to identify genomically close neighbors of NATIVEDY162^T^. The closest bacterial species match was to *Butyrivibrio hungatei* at 96.65% identity (100% coverage), followed by *Butyrivibrio fibrisolvens* at 96.43% identity (99% coverage), and *Butyrivibrio proteoclasticus* at 95.88% identity (100% coverage). None of these values meet the 98.7% similarity threshold required for species identification based on 16S rRNA gene sequences alone [51].

Publicly available genome assemblies for genomically similar species, identified through 16S rRNA gene BLAST, and closely related species based on *Butyrivibrio* phylogenies were compared with NATIVEDY162^T^ to further verify taxonomic placement using Average Nucleotide Identity (ANI). The MUMmer algorithm was first used to generate the alignments for ANI on the basis that this software is adept at aligning large stretches of highly similar sequences and is more stringent than other methods [52-54]. The closest match based on MUMmer ANI was *Clostridium aminophilum* (88.9% identity and 0.555% coverage), however, coverage results to NATIVEDY162^T^ were low (i.e. <6%) in every genome comparison (Table 2). Due to the novelty of NATIVEDY162^T^, it was prudent to compare the alignment results generated by MUMmer with an approach that accounts for more divergent sequences such as BLAST, which compares shorter, local matches [53, 55]. BLAST ANI confirmed *B. proteoclasticus* was the nearest match to NATIVEDY162^T^ at 75.7% identity and 18.7% coverage, followed by *B. fibrisolvens* (75.0% identity/12.9% coverage) and *B. hungatei* (74.9% identity/22.2% coverage; Table 3). Matches to NATIVEDY162^T^ did not meet or exceed a minimum of 95% identity threshold cutoff for ANI, further confirming the NATIVEDY162^T^ identity as a novel species [52-53].

**Table 2.** Whole genome assembly Average Nucleotide Identity (ANI) of NATIVEDY162^T^ compared to genomically similar neighbors and all known *Butyrivibrio* and *Psuedobutyrivibrio* species genome assemblies using MUMmer. All NCBI GenBank accession numbers have been included for reference.

| Microorganism | GenBank Assembly # | ANI (%) | Coverage (%) |
| --- | --- | --- | --- |
| *Clostridium aminophilum* DSM 10710<sup>T</sup> | GCA_000711825 | 88.9 | 0.555 |
| *Pseudobutyrvibrio xylanivorans* DSM 14809<sup>T</sup> | GCA_900141825 | 88.7 | 1.380 |
| *Butyrivibrio proteoclasticus* B316<sup>T</sup> | GCA_000145035 | 88.5 | 5.120 |
| *Butyrivibrio fibrisolven* D1<sup>T</sup> | GCA_037113525 | 87.8 | 2.630 |
| *Coproccoccus comes* ATCC 27758<sup>T</sup> | GCA_025149785 | 86.7 | 0.459 |
| *Butyrivibrio hungatei* MB2003<sup>T</sup> | GCA_001858005 | 86.3 | 4.620 |
| *Pseudobutyrvibrio ruminis* DSM 9787<sup>T</sup> | GCA_900218035 | 85.3 | 0.728 |
| *Roseburia intestinalis* L1-82<sup>T</sup> | GCA_900537995 | 85.2 | 0.439 |
| *Roseburia inulinivorans* DSM 16841<sup>T</sup> | GCA_020731525 | 85.0 | 0.344 |
| *Eshraghiella crossota* DSM 2876<sup>T</sup> | GCA_000156015 | 84.3 | 0.514 |
| *Agathobacter rectalis* VPI 0990<sup>T</sup> | GCA_022453685 | 84.3 | 0.580 |
| *Coproccoccus eutactus* ATCC 27759<sup>T</sup> | GCA_025149915 | 83.5 | 0.544 |

**Table 3.** Whole genome assembly Average Nucleotide Identity (ANI) of NATIVEDY162^T^ compared to genomically similar neighbors and all known *Butyrivibrio* and *Psuedobutyrivibrio* species genome assemblies using BLAST. All NCBI GenBank accession numbers have been included for reference.

| Microorganism | GenBank Assembly # | ANI (%) | Coverage (%) |
| --- | --- | --- | --- |
| *Butyrivibrio proteoclasticus* B316<sup>T</sup> | GCA_000145035 | 75.7 | 18.7 |
| *Butyrivibrio fibrisolvens* D1<sup>T</sup> | GCA_037113525 | 75.0 | 12.9 |
| *Butyrivibrio hungatei* MB2003<sup>T</sup> | GCA_001858005 | 74.9 | 22.2 |
| *Agathobacter rectalis* VPI 0990<sup>T</sup> | GCA_022453685 | 72.9 | 8.54 |
| *Pseudobutyrvibrio xylanivorans* DSM 14809<sup>T</sup> | GCA_900141825 | 72.7 | 10.3 |
| *Coproccoccus eutactus* ATCC 27759<sup>T</sup> | GCA_025149915 | 72.6 | 6.36 |
| *Coproccoccus comes* ATCC 27758<sup>T</sup> | GCA_025149785 | 71.8 | 8.52 |
| *Pseudobutyrvibrio ruminis* DSM 9787<sup>T</sup> | GCA_900218035 | 71.7 | 9.95 |
| *Clostridium aminophilum* DSM 10710<sup>T</sup> | GCA_000711825 | 71.7 | 5.57 |
| *Roseburia inulinivorans* DSM 16841<sup>T</sup> | GCA_020731525 | 71.5 | 7.00 |
| *Roseburia intestinalis* L1-82<sup>T</sup> | GCA_900537995 | 71.4 | 6.58 |
| *Eshraghiella crossota* DSM 2876<sup>T</sup> | GCA_000156015 | 71.3 | 9.60 |

### Phylogeny

A 16S rRNA gene sequence phylogeny was constructed with NATIVEDY162^T^ and all available sequences of the order Clostridiales from the Ribosomal Database Project (RDP). Multiple sequence alignment and nucleotide trimming was performed using MUSCLE aligner and MEGA11 trimming tool, respectively, leaving near full length 16S rRNA sequences [56-57]. A neighbor-joining phylogeny including 649 nucleotide sequences was constructed with 500 bootstrap replicates using MEGA 11 [57]. Annotation and visualization were carried out using FigTree v1.4.4 graphical viewer [58].

NATIVEDY162^T^ placement on the tree was confirmed and 31 sequences were selected for a representative phylogeny that included all known *Butyrivibrio* and *Pseudobutyrivibrio* species, representatives of *Ruminococcus* species, and additional representatives of neighboring *Lachnospiraceae* clades. Nucleotide sequences were aligned using CLUSTALW algorithm and trimmed in MEGA 11 [57, 59]. Bootstrap percent is displayed in decimal form on branch labels calculated from 1000 replicates. The phylogram was rooted using outgroup *Halobacteroides elegans* Z-7287^T^ and all NCBI accession numbers were placed in brackets next to each branch taxon (Figure 1).

**Figure 1.**
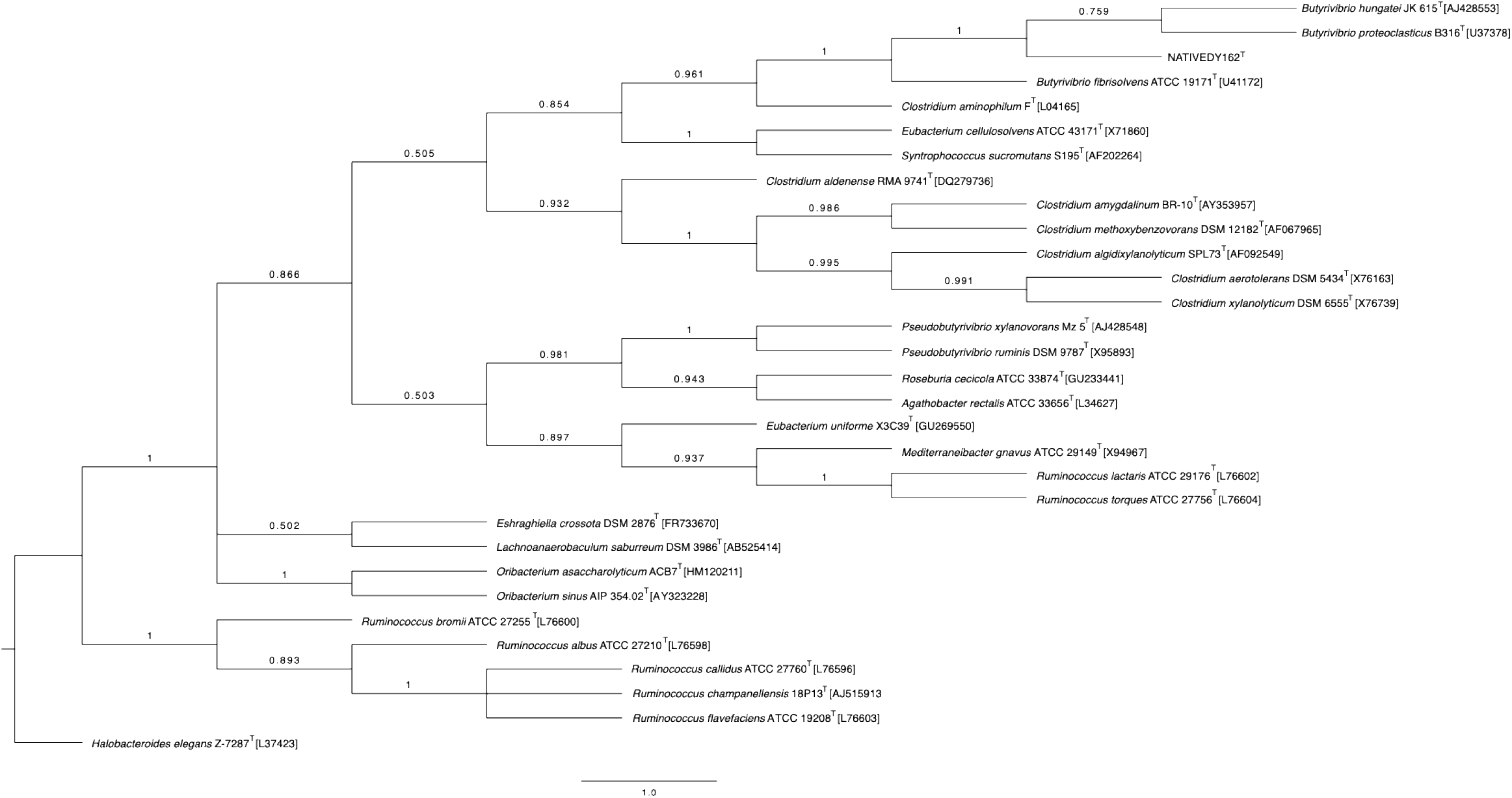
NATIVEDY162^T^ 16S rRNA Gene Phylogram. 16S rRNA gene phylogenetic tree of near full-length gene sequences of NATIVEDY162^T^, *Butyrivibrio* and *Pseudobutyrivibrio* type strains, and close neighbors in the *Lachnospiraceae* family.Phylogeny was inferred using ClustalW multiple sequence aligner, and neighbor-joining tree building model with Mega 11 software. Tree visualization was performed using FigTree and outgroup rooting with *Halobacteroides elegans* Z-7287^T^. Branches include bootstrap values in decimal form calculated from 500 bootstrap replicates. All NCBI GenBank accession numbers are available in brackets listed after taxa. NATIVEDY162^T^ and all *Butryrivibrio* species can be seen at the top branches of the tree.

The closest neighbors were determined to be *B. hungatei, B. proteoclasticus*, and *B. fibrisolvens*, which is in accordance with the results noted in the BLAST ANI output and the phylogenetic clustering noted in Palevich et al. 2019 [31]. NATIVEDY162^T^ was also placed near members previously found to be related to and classified as the *Clostridium* subphylum XIVa, including *Clostridium aminophilum, Eubacterium cellulosolvens, Syntrophococcus sucromutans*; a phylogenetic characteristic typical of *Butyrivibrio* species (18, 24, 27, 29, 60].

### Physiology and Chemotaxonomy

pH tolerance was confirmed by growing NATIVEDY162^T^ in a modified tryptic soy broth medium, which consisted of 3.0 g/L casein (MP Biomedicals, Irvine, CA), 3.0 g/L peptone from soybean (Sigma-Aldrich, St. Louis, MO), 2.5 g/L glucose (Nutricost, North Vineyard, UT), 5.0 g/L NaCl (Spectrum, New Brunswick, NJ), 2.5 g/L dipotassium phosphate (Spectrum, New Brunswick, NJ) and 8.0 g/L Sensiferm 605 yeast extract (Sensient, Hoffman Estates, IL) in deionized water. Media was aliquoted into Balch tubes in 20 mL volumes and sparged at 80% N_2_ / 20% CO_2_ [v/v] for 20 minutes before capping with butyl septums and aluminum crimp seals. All media was autoclaved for 25 mins at 121°C / 15 psi and allowed to cool prior to adding the following sterile additions to each tube: 1X Wolfe’s mineral solution and 0.5 g/L cysteine-HCl (Spectrum, New Brunswick, NJ). A series of tubes was pH adjusted in 0.5-unit increments ranging from pH 4.0 to pH 9.0 with either sterile 0.5M hydrochloric acid or 0.5M sodium hydroxide. Each pH condition was inoculated with 1.0% [v/v] NATIVEDY162^T^ and incubated at 37°C. Growth was observed and tracked over a period of 65 hours by optical density 600nm (OD_600_) readings. NATIVEDY162^T^ did not grow in the pH range of 8.0-9.0, grew very poorly from pH 4.0-5.0, and grew slowly and at mid-range at pH 5.5-6.0. It preferred the pH range from 6.5-7.5 and yielded the highest titers at 7.0.

Catalase, oxidase, and sulfur-indole-motility (SIM) tests were performed to confirm additional biochemical characteristics of NATIVEDY162^T^. SIM differential media was created based on a 2009 protocol by MacWilliams [61]. The medium recipe included 30 g/L Difco tryptic soy broth (BD Biosciences, Franklin Lakes, NJ), 0.5 g/L cysteine-HCl (Spectrum, New Brunswick, NJ), 3.0 g/L agar (Spectrum, New Brunswick, NJ), 0.2 g/L ammonium iron(III) citrate (Sigma-Aldrich, St. Louis, MO), 0.02 g/L sodium thiosulfate (Macron, Radnor, PA), and 5.0 g/L tryptophan (Sigma-Aldrich, St. Louis, MO) in deionized water. Media was aliquoted into test tubes, autoclaved for 25 mins at 121°C / 15 psi, and was placed in an anaerobic chamber (Bactron, Cornelius, OR) for 72 hours prior to stab inoculation with a needle. Results after 48 hours indicated NATIVEDY162^T^ does not convert sulfur into hydrogen sulfide, does not produce indole from the breakdown of tryptophan, and is non-motile. Catalase presence or absence was tested using a 3% (v/v) hydrogen peroxide solution and oxidase capabilities were tested using a 1.2% tetra-methyl-p-phenylenediamine solution [62-63]. NATIVEDY162^T^ is indole-, oxidase-, and catalase-negative, which is characteristic of other *Butryrivibrio* species [28].

Carbohydrate utilization capabilities were determined using the API 50CH carbon panel (BioMérieux, Marcy-l’Étoile, France). Cells were first grown to late log phase using the modified TSB medium recipe at pH 7.0 as described above. Cells were harvested by centrifugation at 16,000 x g for 3 mins and culture supernatant was removed by micropipette. The cell pellet was resuspended in a carbohydrate-free tryptic soy broth medium and inoculated into API test strip wells. Test strips were incubated in anaerobic conditions for 48 hours before adding 5 µL 0.85% bromothymol blue dye indicator to confirm presence or absence of acid production. NATIVEDY162^T^ was confirmed to utilize D-glucose, L-rhamnose, esculin/ferric citrate, D-lactose, D-trehalose and grew weakly in L-arabinose, D-xylose, D-galactose, D-mannose, D-cellobiose, D-maltose, starch, and glycogen. It did not grow in any of the other carbohydrates tested. The full list of carbohydrates can be found in Supplementary Table 1.

Starch degradation ability was additionally verified using the protocol detailed by Lal & Cheeptham [64]. Growth medium chosen for the assay was a custom reinforced clostridial agar medium (RCA) constructed by combining 10.0 g/L peptone (Research Products International, Mount Prospect, IL), 10.0 g/L beef extract (Sigma-Aldrich, St. Louis, MO), 3.0 g/L yeast extract (Sigma-Aldrich, St. Louis, MO), 5.0 g/L NaCl (Spectrum, New Brunswick, NJ),3.0 g/L sodium acetate (Sigma-Aldrich, St. Louis, MO), and 12.0 g/L agar (Spectrum, New Brunswick, NJ) in deionized water. Three different test starch substrates (i.e. soluble starch, starch from corn, or amylose from potato; Sigma-Aldrich, St. Louis, MO) were added to three separate batches of media at 1.0 g/L and autoclaved at 121°C / 15 psi. Plates were maintained in an anaerobic chamber (Bactron, Cornelius, OR) for 72 hours prior to inoculation. NATIVEDY162^T^ was streaked onto all three media types and incubated anaerobically at 37°C for 72 hours. Iodine stain post bacterial growth confirmed NATIVEDY162^T^ was positive for hydrolysis of soluble starch, starch from corn, and amylose from potato (Supplementary Figure 3).

The protocol by Szydlowski et al., 2015 was adapted to corroborate cellulose digestion by NATIVEDY162^T^ [65]. Custom RCA medium lacking carbohydrates or starch as described above was additionally modified with 0.2 g/L Congo red powder and 1.0 g/L either cellulose powder or carboxymethyl cellulose sodium salt (CMC) (Sigma-Aldrich, St. Louis, MO) to test both insoluble and soluble forms of cellulose. Media was autoclaved at 121°C / 15 psi and poured plates were placed in an anaerobic chamber (Bactron, Cornelius, OR) for 72 hours prior to inoculation. NATIVEDY162^T^ was applied to both media types with an inoculating loop and incubated anaerobically at 37°C for 72 hours. Photos were captured with a Scan Interscience 500 (Interscience, Woburn, MA) with a backlighting filter. NATIVEDY162^T^ was confirmed as positive for the cellulolytic phenotype of both CMC and cellulose (Supplementary Figure 4).

Metabolites were measured by High Performance Liquid Chromatography (HPLC) to confirm the fermentation byproducts of NATIVEDY162^T^. Cells were first grown anaerobically to late log phase in modified anaerobic TSB medium as previously described. A 1 mL sample was removed, centrifuged at 16,000 x g for 3 mins, and supernatant collected and filtered through a 0.22 µm polyestersulfone membrane. Samples were analyzed using an Agilent 1260 Infinity II System equipped with a refractive index detector (Agilent, Santa Clara, CA). Separation was achieved on a Hi-Plex H column at 35°C with 5 mM sulfuric acid used as the mobile phase at a flow rate of 0.2 mL/min. Fermentation of glucose by NATIVEDY162^T^ resulted in the production of lactate, butyrate, and minor amounts of acetate. Table 1 shows a comparison of known carbohydrate utilization and metabolite production capabilities by known *Butyrivibrio* and *Pseudobutyrivibrio* species to that of NATIVEDY162^T^. Cellulolytic capabilities may serve as a distinguishing feature of NATIVEDY162^T^, since most *Butyrivibrio* have been reported to be incapable of cellulose degradation [31].

### Nomenclature and Description of *Butyrivibrio azoria* sp. nov

The cultural landscape of the dairy industry in the United States, and specifically in California, is rich in Portuguese influence [66-67]. Portuguese migrants began to settle in California’s agriculturally abundant San Joaquin Valley during the time of the 1848 Gold Rush [66, 68-69]. Communities formed in Hanford and Tulare counties into the early 1900s as Portuguese immigrants sought economic opportunities in the dairy, fishing, timber, and sheep herding industries [66, 68-70]. By the 1930s, people of Portuguese descent represented the numerical majority in the California dairy industry [67, 69]. NATIVEDY162^T^ was isolated from the rumen of a dairy cow located in the San Joaquin Valley. Azorean Portuguese people that immigrated to California had major impacts on the dairy industry with almost 75% diary ownership and still made up almost half of the dairy owners in California into the early 2000s [70-71]. It is because of the history and impact on the dairy industry in California we therefore propose the name *Butyrivibrio azoria*, which underscores the importance of the contributions of the Azorean and Portuguese dairy farmers.

*Butyrivibrio azoria* (azoria L. m. gen. n. place/land of Azores)

*Butyrivibrio azoria* type strain NATIVEDY162^T^ is a novel obligately anaerobic, slightly curved rod (0.550-0.7 μm x 2.5-3.6 μm) that forms singlets or short chains. It stains Gram negative, is non-motile, and is indole, oxidase, and catalase negative. When cultured on unmodified TSB agar solid media, colonies appear as opaque white, small, and circular with smooth edges. *B. azoria* NATIVEDY162^T^ is cellulolytic, can hydrolyze starches, and growth is also supported by utilize D-glucose, L-rhamnose, esculin/ferric citrate, D-lactose, D-trehalose.

When grown in a custom tryptic soy broth it clumps to form aggregates and converts glucose to lactate and butyrate, with a minor amount of acetate production.

*B. azoria* NATIVEDY162^T^ has two chromosomes: one main chromosome, as well as a chromid which contains genes that support its cellulolytic and amylolytic phenotype (i.e. xylanases, endoglucanase, cellulase, glycosyl hydrolase). Chromosome one is 3,687,363 bp and 40.42 mol% G-C content and the chromid is 449,257 bp and has DNA G-C content at 39.81 mol%. A 16S rRNA gene phylogram supports NATIVEDY162^T^ clusters with other *Butyrivibrio* strains and ANI analysis shows its closest neighbor of a known species being *B. proteoclasticus*.

The type strain NATIVEDY162^T^ (= JL13D10^T^ = NRRL B-68525^T^) was isolated from rumen content of a healthy Holstein dairy cow located in a dairy farm in Corcoran, CA, USA.

### Protologue

All sequences were deposited in the NCBI GenBank.

NATIVEDY162^T^ [= JL13D10^T^] 16S rRNA Accession NCBI GenBank # PQ046053

NATIVEDY162^T^ [= JL13D10^T^] WGS Accession NCBI GenBank # CP172573 & CP172574

Strain Deposit and Availability: =NRRL B-68525^T^

## AUTHOR STATEMENTS

### Ethical Statement

Sampling procedures were approved by veterinarians and the IACUC at Native Microbials, Inc.

## ABBREVIATIONS

ANI: Average nucleotide identity
CAZymes: carbohydrate active enzymes
CMC: carboyxymethyl cellulose sodium salt
ONT: Oxford Nanopore Technologies
RCA: reinforced clostridial agar
RDP: Ribosomal Database Project
TSB: Tryptic Soy Broth

