## Supplementary Materials for "*Butyrivibrio azoria* sp. nov., a novel cellulolytic microorganism isolated from the rumen of a Holstein dairy cow"

**Supplementary Figure 1. Methylene Blue Stain**

Methylene blue stain of NATIVEDY162$^{T}$ visualized under 1000X magnification. Cells were grown anaerobically in TSB medium for 72 hours until reaching late log phase.

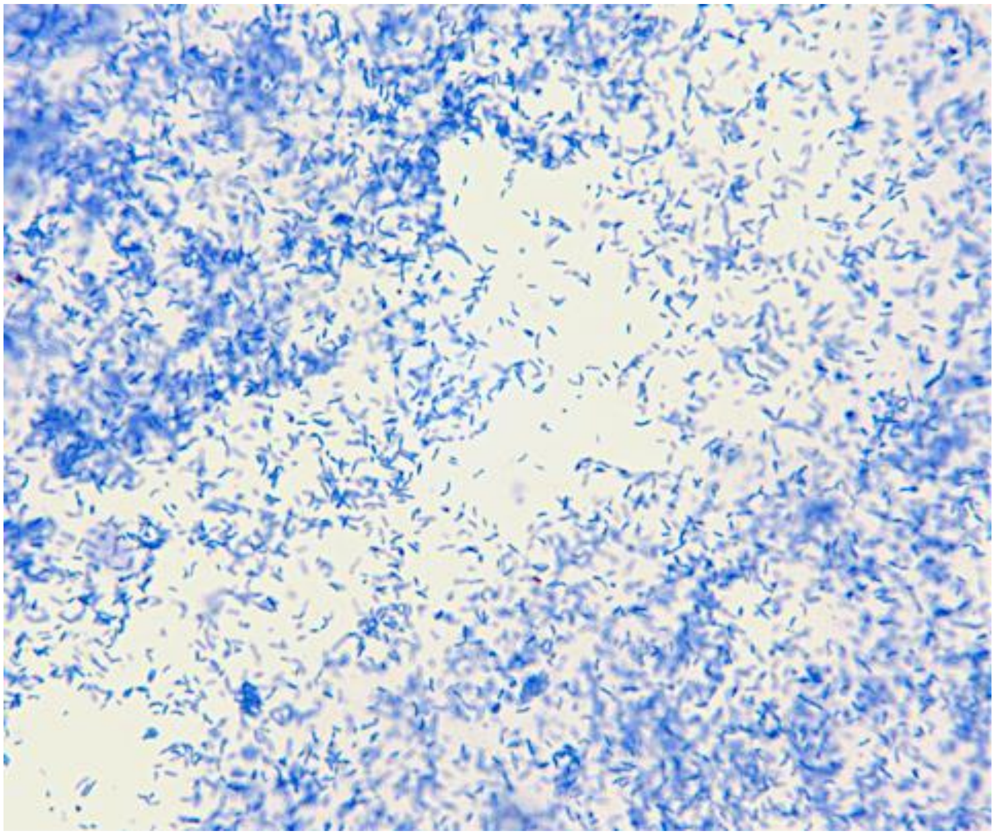

**Supplementary Figure 2. Gram Stain**

Gram stain of NATIVEDY162$^{T}$ visualized under 1000X magnification. Cells were grown anaerobically in TSB medium for 72 hours until reaching late log phase.

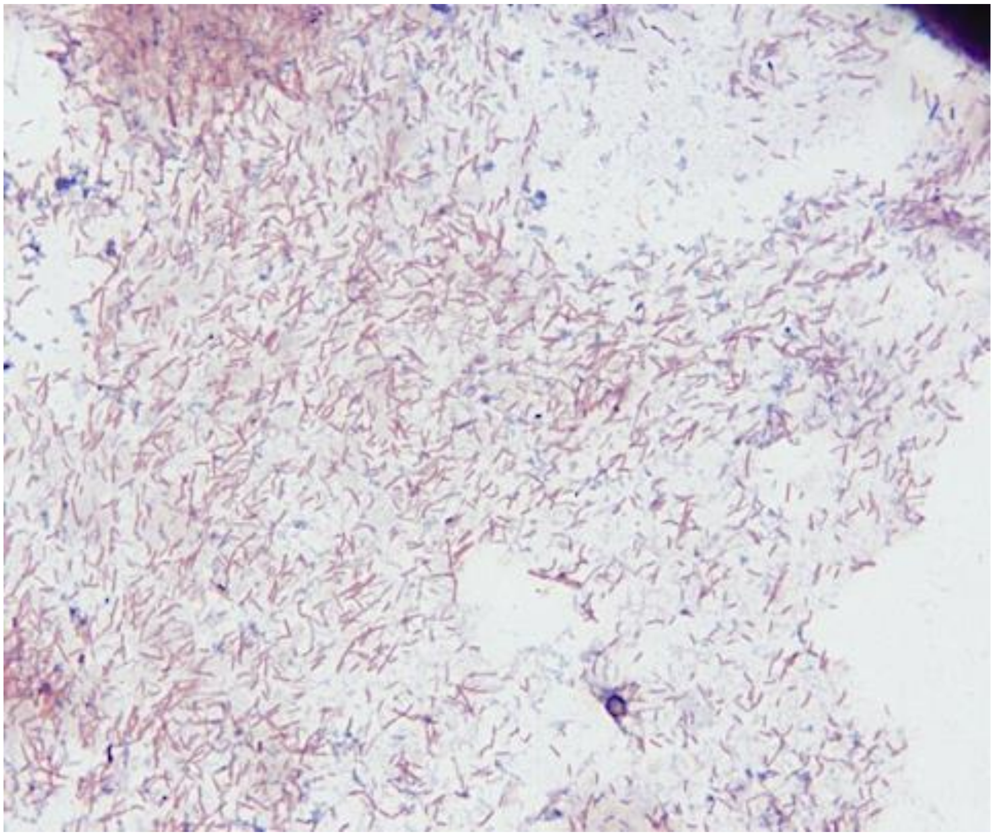

**Supplementary Table 1. API 50CH for NATIVEDY162$^{T}$**

Carbohydrate utilization panel for NATIVEDY162$^{T}$ using BioMérieux's API 50CH carbon test strips. Minus or plus signs indicate a negative or positive result, while a symbol of 'w' is indicative of weak growth.

| Component | NATIVE DY162$^{T}$ Growth (+/-) | Component | NATIVE DY162$^{T}$ Growth (+/-) |
|---|---|---|---|
| Control | - | Esculin/Ferric Citrate | + |
| Glycerol | - | Salicin | - |
| Erythritol | - | D-Cellobiose | w |
| D-Arabinose | - | D-Maltose | w |
| L-Arabinose | w | D-Lactose | + |
| D-Ribose | - | D-Melibiose | - |
| D-Xylose | w | D-Saccharose | - |
| L-Xylose | - | D-Trehalose | + |
| D-Adonitol | - | Inulin | - |
| Methyl-BD-xylopyranoside | - | D-Melezitose | - |
| D-Galactose | w | D-Raffinose | - |
| D-Glucose | + | Starch | w |
| D-Fructose | - | Glycogen | w |
| D-Mannose | w | Xylitol | - |
| L-Sorbose | - | Gentiobiose | - |
| L-Rhamnose | + | D-Turanose | - |
| Dulcitol | - | D-Lyxose | - |
| Inositol | - | D-Tagatose | - |
| D-Mannitol | - | D-Fucose | - |
| D-Sorbitol | - | L-Fucose | - |
| Methyl-aD-Mannopyranoside | - | D-Arabitol | - |
| Methyl-aD-Glucopyranoside | - | L-Arabitol | - |
| N-AcetylGlucosamine | - | Potassium Gluconate | - |
| Amygdalin | - | Potassium 2-KetoGluconate | - |

**Supplementary Figure 3. NATIVEDY162$^{T}$ Streaked Out on Starch Media**

Starch hydrolysis assay photos captured in natural light. NATIVEDY162$^{T}$ was streaked on three starch substrate solid media: soluble starch (left), starch from corn (middle), and amylose from potato (right). Plates were incubated in anaerobic conditions for 72 hours at 37°C. Top row photos show unstained plates with microbial growth, while bottom row photos have been stained with iodine. A clearing/halo surrounding microbial growth within the purple iodine stain is indicative of a positive result.

Soluble Starch | Starch from Corn | Amylose from Potato

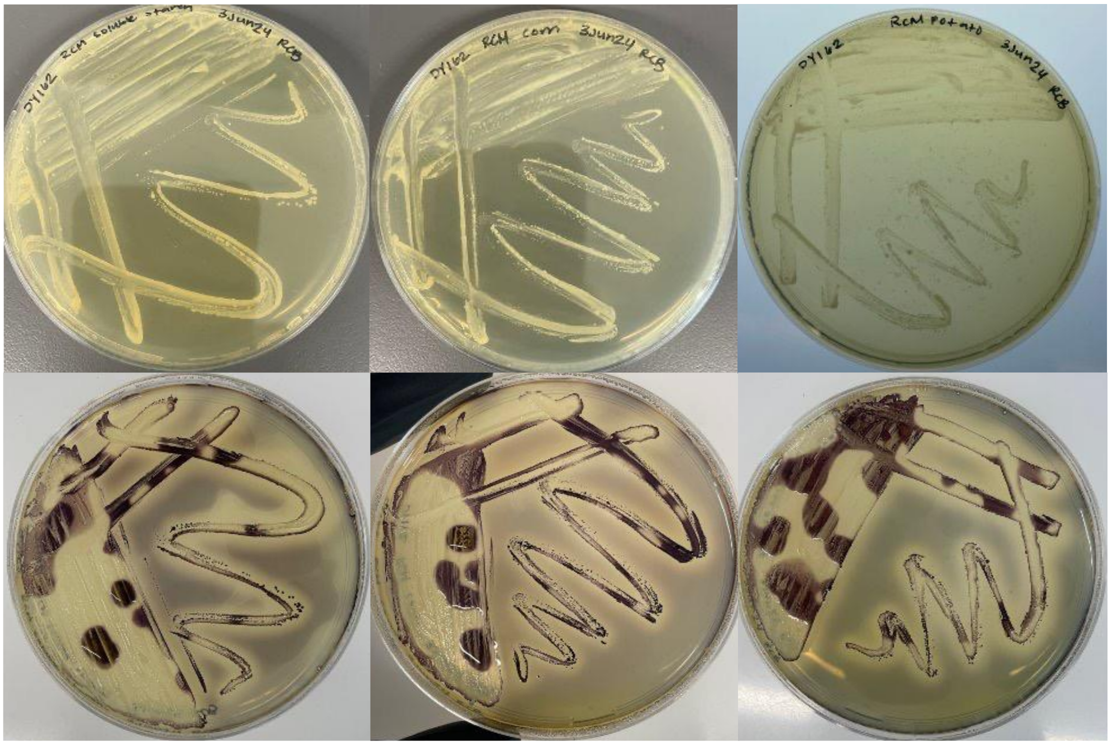

**Supplementary Figure 4. NATIVEDY162$^{T}$ Grown on Cellulose and CMC Congo Red Media**

Cellulose digestion assay photos of NATIVEDY162$^{T}$ with backlighting (right) and in natural lighting (middle). Uninoculated plates are pictured left. NATIVEDY162$^{T}$ was grown in anaerobic conditions for 72 hours at 37°C on custom RCM CMC medium (top row) and cellulose medium (bottom row). A clearing of the Congo red dye (usually with yellow color change) surrounding microbial growth indicates cellulose or CMC digestion.

| Uninoculated | Inoculated | Inoculated |
|---|---|---|

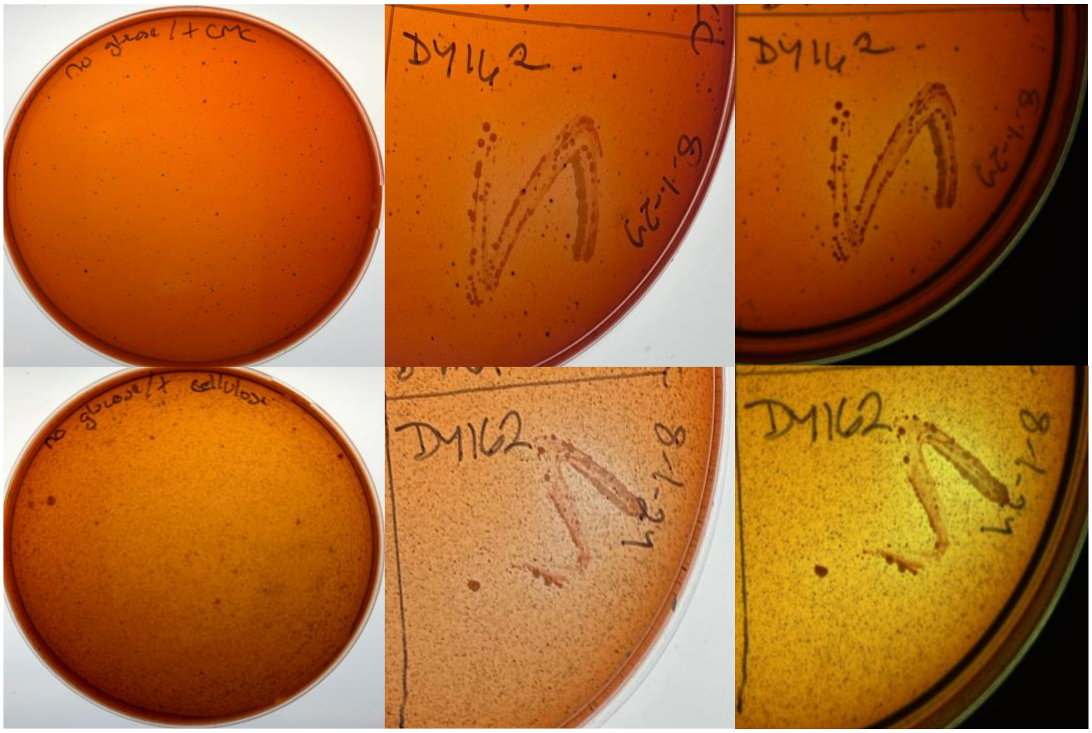
